# Investigating Effects of Outcome Controllability and Error Attribution on Proactive Attentional Control: Insights from EEG and Cognitive Modelling

**DOI:** 10.64898/2026.03.03.709239

**Authors:** Luisa A. Grote, Daniel Schneider, Edmund Wascher, Stefan Arnau

## Abstract

Sense of agency (SoA), the experience of controlling one’s actions and their consequences, is crucial for self-representation and adaptive goal-directed behavior. Classic comparator models explain SoA as the match between predicted and actual sensorimotor outcomes, whereas inference-based and Bayesian accounts emphasize cue integration and probabilistic weighting. Besides the influence of action-outcome contingencies on SoA, the feedback effect of perceived SoA on cognitive processing is also crucial for cognitive performance. Much of today’s cognitive work is performed through interaction with devices that are not entirely reliable or are prone to operator error. Against this background, it is of particular interest whether the impact of an expectancy violation differs depending on whether the outcome is attributed to a malfunctioning system or to one’s own mistake. To investigate this, the present EEG study deploys manipulated performance feedback in a color-discrimination task, while EEG was recorded. Thirty-five participants performed in this task with periods of veridical feedback, periods with feedback simulating an increased error rate, and periods of feedback suggesting malfunctioning response buttons. Behavioral performance was decomposed using the EZ-diffusion model, and time-frequency EEG analyses focused on event-related alpha, beta, and theta oscillations. The participants responded significantly slower in the self-attribution of errors condition compared to neutral feedback, and also significantly slower in the system-attribution of errors condition compared to self-attribution of errors. Decomposing behavior using drift-diffusion modeling indicates that a general decrease of response times with manipulated feedback can be attributed to decreased drift rates, whereas the difference between the self and system error conditions are reflected in the non-decision time. In the EEG, the manipulated feedback was reflected in attenuated decreases of occipital alpha and sensorimotor beta power during the cue-target interval. Furthermore, system-versus self-attributed errors elicited stronger feedback-locked midfrontal theta responses. Our findings suggest a functional dissociation within the agency inference process, where perceived controllability regulates preparatory investment of cognitive resources, while the attribution of action-outcome discrepancies seem to modulate sensory processes or motor-execution.

## Introduction

Sense of agency (SoA), the feeling of being in control of one’s actions and their effects on the environment (Haggard, 2017; Gallagher, 2000), is central to goal-directed behavior (Chambon et al., 2014; Kaiser et al., 2021). It provides a basis for self-representation and mediates the adaptive use of external feedback to guide and control (Kaiser et al., 2021; Synofzik et al., 2013; Gentsch & Synofzik, 2014). Disturbances of SoA have been linked to psychiatric disorders such as schizophrenia (Carruthers, 2012; Moore & Fletcher, 2012; Bayne, 2008), underscoring its importance for healthy cognition and behavior (Gallagher, 2013; Haggard, 2017; Verschoor & Hommel, 2017). There is emerging evidence that SoA constitutes a cognitive state that in return affects cognitive processing of a task (Ren et al, 2023). The present study investigates how a manipulation of SoA might affect task engagement and furthermore investigates the role of error attribution.

SoA has traditionally been addressed by the comparator model, which has been adapted from motor control literature (Wolpert et al., 1995). The comparator model leans on the assumption that the brain uses forward models to predict the sensory consequences of an action (Blakemore et al., 2000) using an efference copy of motor commands (Feinberg, 1978; Haggard, 2017; Wen & Haggard, 2020). If the outcome matches the predictions, it is inferred to be self-caused, while incongruent feedback is externally attributed (Frith, 1987; Gallagher, 2000). While fundamental for most accounts for SoA, this approach focusses narrowly on sensorimotor prediction.

Inference-based accounts expand this view, emphasizing the integration of sensorimotor evidence with contextual priors for the establishment of SoA (Synofzik et al., 2008; Moore & Fletcher, 2012; Frith 2012). They propose that SoA does not arise from a single prediction-outcome comparison, but from the integration of multiple cues with varying informational value, such as differing reliability (Synofzik et al., 2008). These cues may include motor predictions derived from efference copies, as proposed by the comparator model, but also proprioceptive and visual feedback, contextual information and higher-order beliefs about one’s own control. For example, Gentsch et al. (2012) demonstrated that SoA depends on the integration of both motor and non-motor anticipatory signals, with their relative contribution depending on cue reliability. In this framework, SoA is conceptualized as the outcome of a reliability-weighted cue integration process rather than a single comparator operation (Moore & Fletcher, 2012). This implies that changes in perceived feedback reliability may shift the weighting of predictive and contextual cues, thereby altering the perceived SoA.

More recently, computational models rooted in Bayesian theory have formalised SoA as the result from a probabilistic inferencing process (Legaspi & Toyoizumi, 2019; Dutta, 2025; Tanaka, 2024). These accounts integrate comparator-derived prediction errors, contextual information, intentions and social cues through multilevel probabilistic computations, weighted according to their reliability. Such computational frameworks therefore capture both sensorimotor and cognitive influences on SoA. Consistent with this view, Ren et al. (2023) showed that the response tendencies in a Go/Nogo-Task are modulated by local SoA manipulations. They found a higher SoA to be associated with faster responses in Go-trials, whereas a lower SoA was associated with a decreased accuracy in Nogo-trials. Such results clearly demonstrate that SoA is not only a consequence of perceived action-outcome contingencies, but that it also constitutes a cognitive state that in return affects cognitive processing and task engagement.

A modulatory effect of SoA and task engagement can also be aligned with recent frameworks on the allocation of cognitive resources. It is hypothesized that the allocation of cognitive effort is based on the expected reward of successful task performance (Shenhav et al., 2013). It has also been shown that the expenditure of cognitive resources is modulated by its expected efficacy. Frömer et al. (2021) contrasted an experimental condition in which the participants were rewarded for performing good enough, to a condition where the reward scaled linearly with performance. Cognitive effort was significantly higher in the latter condition, showing that individuals put effort into a task when it is more likely to translate into reward.

A related question that is also insufficiently explored concerns the attribution of errors. The allocation of cognitive resources regarding SoA does not necessarily depend on the perceived agency alone, but also on the perceived reason for failure. When an intended action-outcome is not met, we must infer whether this deviation stems from one’s own performance error within an intact action-outcome contingency, or from a disruption of the causal mapping itself (Desantis et al., 2012; Synofzik et al., 2008). The attributional inference determines whether the beliefs about one’s own performance and outcome-controllability are updated (Tanaka, 2024; Legaspi & Toyoizumi, 2019; Friston, 2012). Failing to achieve an intended action goal may influence subsequent actions and control allocation in different ways, depending on whether the failure is attributed to one’s own mistake or to an external cause (Ullsperger et al., 2014; Cavanagh & Frank, 2014; Taylor et al., 2007).

To address these questions, the present EEG study deployed specific performance feedback conditions to manipulate SoA in a calibrated colour discrimination task. Three within-subject conditions were contrasted. First, a veridical feedback condition provided accurate feedback by displaying the points gained or lost on each trial based on whether the participant’s response was correct or incorrect. Second, a self-attribution of errors condition was implemented, in which 33% of correctly answered trials were falsely presented as errors. In this condition, the feedback indicated that the participant had selected the chosen color, but that this response was incorrect, even though it was in fact correct. Third, a system-attribution of errors condition was included, in which 33% of correctly answered trials were also falsely presented as errors. However, in this case the feedback indicated that the alternative color, rather than the one selected by the participant, had been chosen, and that this response was incorrect. Thus, participants should have had the impression that they had responded correctly, but that their response had been compromised by the recording software. As an additional experimental factor, we also manipulated reward for successful performance on a trial-to-trial basis to investigate whether reward-related expenditure of cognitive effort interacts with SoA-related effects.

We hypothesised that the perceived decrease in outcome controllability inferred from increased error rates in self- and system-attributed error conditions would modulate the participants’ cognitive state, resulting in the withdrawal of cognitive effort when action-outcomes cannot be controlled as anticipated. To test this, we combined behavioral measures of response time and accuracy with drift diffusion modelling (DDM) using the EZ-diffusion model (Wagenmakers et al., 2007). Furthermore, event-related EEG data was analysed using time-frequency decomposition to investigate cue- and target-related oscillatory activity on the one hand, and response- and feedback-related activity on the other hand.

Regarding behavior, we expect that the reduced perceived controllability in the manipulated feedback conditions impair task performance by affecting cognitive engagement, leading to increased response times. A reduced task engagement is often associated with down-regulated proactive cognitive control processes (c.f. Braver et al. 2012). We therefore hypothesize that DDM modelling of response times would reveal reduced drift rates in unreliable feedback conditions relative to veridical feedback. Moreover, we expect system-attributed errors to affect performance even more compared to self-attributed errors, as the action-outcome conflict is more salient and perceived as clearly outside of the participants’ controllability.

Our experimental design allows us to use the EEG to observe proactive control allocation and performance monitoring processes independently. Proactive control has been linked to modulations of ongoing oscillatory activity. Occipital alpha power and beta power over sensorimotor areas prior to the target onset have been associated with visual attention and motor readiness, respectively (Jensen & Mazaheri, 2010; Klimesch, 2012; Foxe & Snyder, 2011). We therefore hypothesize that occipital alpha power as well as central beta power (Engel & Fries, 2010) during the cue-target interval are likely to be modulated by the experimental manipulations. We expect that reliable performance feedback would be associated with a stronger decrease of alpha and beta power compared to self-and system attributed error contexts. Regarding the attribution of errors, we expect to observe specific differences in EEG measures reflecting monitoring processes. Here, we focus our analyses on feedback-related theta activity at mid-frontal recording sites. Midfrontal theta power has been described as a measure reflecting the need for control, reflecting error- monitoring and feedback processing (Cavanagh & Frank, 2014, Kaiser et al., 2021; Luft et al., 2013).

Such findings would indicate that perceived outcome-controllability shapes proactive control allocation, whereas differential processing of self-and system-attributed errors reflects evaluative monitoring processes. This would suggest that distinct components of agency, namely the preparation of outcome predictions, and the monitoring of actual action-outcome contingencies, would operate at different stages of cognitive processing.

## Methods

### Participants

Overall, the data of 38 participants were collected. The inclusion criteria were right-handedness, intact colour-vision, no suffering from any psychiatric or neurological disorder, as well as being in the age range of 18 to 30. Handedness was measured using the Edinburgh Handedness Inventory (Oldfield, 1971), colour vision was tested using the Ishihara test (reference?). The data of 2 participants had to be excluded from further analysis due to poor data quality, 1 participant did not finish the experiment. According to an a priori power analysis using *G*power 3.1 (Faul et al., 2007), the remaining sample of 35 participants (13M/21F/1 diverse; age M=24.63, SD=4.66) is sufficient to detect medium sized effects with a power of .95. The study was approved by the local Ethics Committee of the Leibniz Institute for Working Environment and Human Factors. Written informed consent was obtained prior to starting the experiment. Participants were compensated with 13€/hour and an additional performance-based reward up to 20€.

### Procedure

Upon arrival, participants were informed about the proceedings of the experiment, the monetary compensation, and data protection, after which they signed a consent form and filled out questionnaires about their demographic background. Following, they were positioned in the EEG chamber for the preparation of the EEG cap, during which they filled out several psychometric questionnaires (Boredom Proneness Scale, Internal-External Locus of Control Short Scale (IE-4), Beck Depression Inventory, General Self-Efficacy Scale).

After EEG cap preparation, participants completed a 10-minute calibration phase consisting of 120 trials, which also served as a familiarisation phase for the participants. Stimuli were displayed on a 20” monitor with a refresh rate of 100Hz located at 1.3 meters distance from the participant. Calibration trials consisted of: (1) a fixation point, (2) a dichromatic pixel field, consisting of 23 by 23 pixels (≤1000ms; orange #F9C826/cyan #27EDD0), and (3) a button-press response (left/right index finger) indicating the dominant colour, followed by (4) veridical feedback (correct colour + lateralized smiley). Calibration grids presented a continuous range of colour ratios (40–49%), generated by linearly interpolating between the bounds so that each trial contained a unique ratio. Because these ratios were then randomized in order, participants experienced a smooth, stepless manipulation of difficulty rather than categorical difficulty levels. Calibration trials were then analysed using separate logistic regressions for blue- and yellow-majority grids, modelling accuracy as a function of symmetric difficulty (|ratio–0.5|). For each colour, a 80% performance threshold was estimated by inverting each regression function and converting the resulting difficulty value back into the corresponding pixel ratio. These participant-specific thresholds of each colour defined the individualized stimulus ratios used during the main experiment to rule out perceptual biases. We selected the 80% threshold to ensure sufficient task competence to establish SoA over the task by feeling in control of action-outcomes, while preserving some uncertainty necessary to increase the perceived error rate in the self-attribution of errors condition without immediate notice of the experimental manipulation.

The main task operationalizes a 3 (veridical feedback, self-attribution of errors, system attribution of errors) × 2 (high vs low reward) factorial design. The factor reward was manipulated on a trial-to-trial basis, the factor feedback type in a block-wise manner. The structure of the trials is depicted in Figure 1. Each trial began with a fixation point, followed by a reward cue, displayed for 500ms. Counterbalanced across participants, either a circle or a diamond indicated high- (10 points) and low-reward trials (1 point). After a 1000ms inter-stimulus-interval (ISI), a dichromatic pixel field was presented until a response was given or a maximum response time of 1000ms was reached. It was the task of the participants to indicate the dominant colour by pressing the left or right response button with the respective index finger. Time-locked to the response, condition-specific feedback was presented for 500ms. The feedback informed about the selected colour, that is, the button pressed by the participant, and the number of points that were obtained or missed out on according to the amount obtainable as previously indicated by the reward cue.

**Figure 1.**
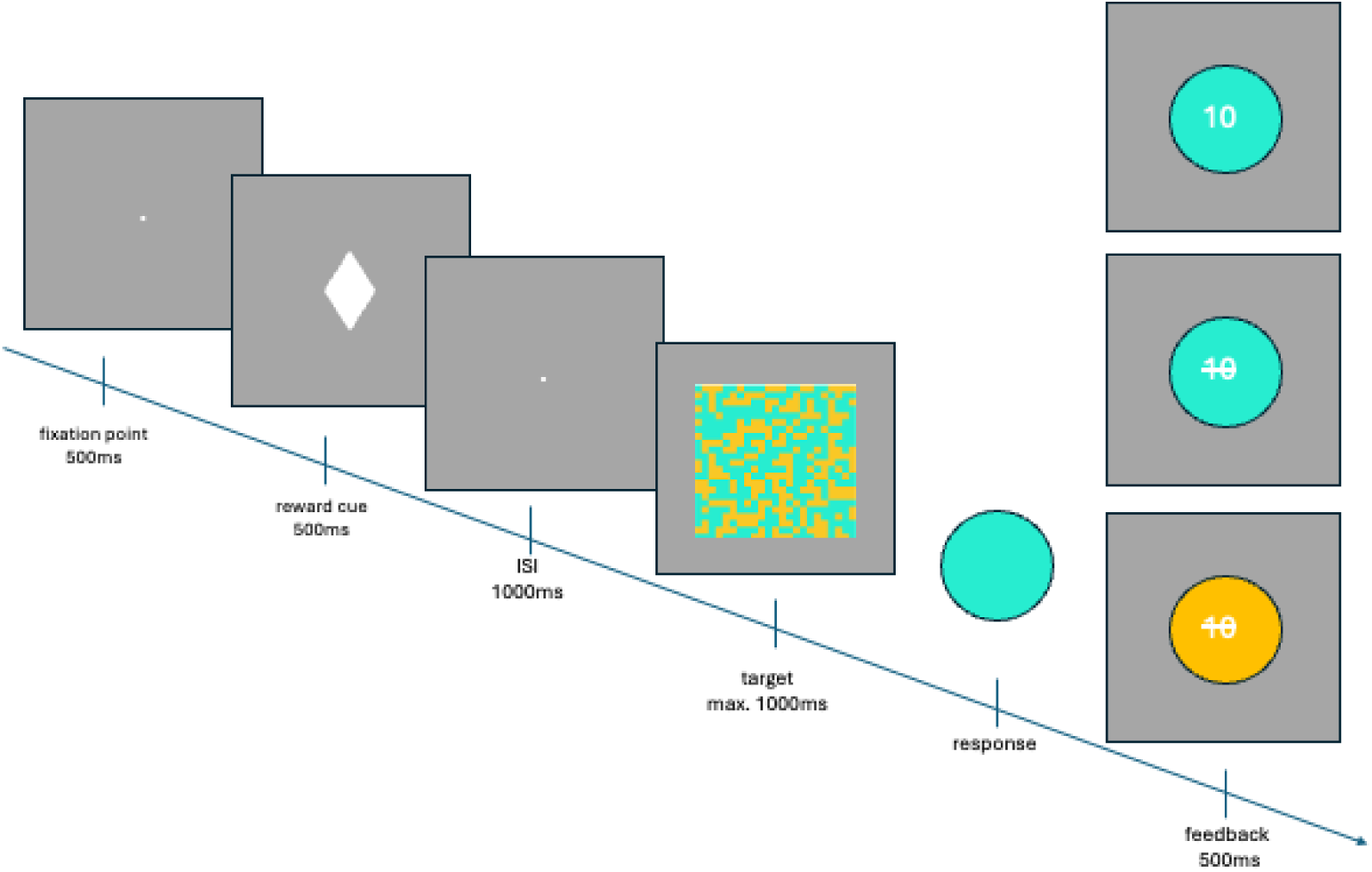
Time course of stimulus presentation. Each trial started with a fixation point, followed by a reward cue shown for 500ms indicating high or low reward trials. After a 1000ms inter-stimulus-interval, the target was presented for a maximum duration of 1000ms, during which the participants had to indicate the dominant colour with a button press of their left or right index finger. Feedback was presented for 500ms immediately upon button press, consisting of a circle in the colour of the button that was pressed (Veridical feedback) and the amount of points gained or missed out on, the colour pressed with manipulated feedback indicating missing out of points due to a wrong answer (self-attributed error) or the opposite colour of which was pressed and missed out points due to a wrong answer (system-attributed error).

In the veridical feedback condition, this feedback always corresponded to the given responses. In the self-attribution of errors condition, the feedback of each correctly answered trial had a 33% chance of being flipped to indicating that no points were obtained. This means that the feedback showed the colour that was pressed by the participant, but the points were crossed out, suggesting a wrong answer. We hypothesize that in this condition, the participants would attribute the failure to obtain the points to their own mistake. Due to the a priori calibration of the task to an accuracy of 0.8, participants should not have maximum certainty about the correctness of their answer when they give it, but still a sufficient degree of task competence to feel in control of the outcome and, respectively, accountable for the mistake. In the system-attribution of errors condition feedback, each correctly answered trial also had a 33% chance of being flipped to an allegedly wrong answer. This time, the feedback was not only related to the points obtained, but it was also indicated that the participants had pressed the wrong button (see Figure 1). Here, the manipulation should be very salient to the participants, as the response-mapping to the left and right buttons was kept constant for each participant, and even the physical buttons themselves are in the respective colour. Moreover, participants had received information prior to the start of the experiment that the computer would sometimes record their answers incorrectly. We therefore hypothesize that the participants would attribute the failure to obtain the points in this condition to an error in the system. Because feedback conditions were manipulated block wise, there were 12 blocks total, each consisting of 120 trials. Each participant saw the veridical feedback condition first. Subsequent blocks were randomized without repetitions. Participants took short breaks every two blocks and a longer break after 6 blocks. The total duration of the experiment was approximately 2.5h.

Throughout the experiment, thought probes were collected every 10–50 trials (every 30 trials on average). These thought probes assessed task-focus (How well can you focus?), performance satisfaction (How satisfied are you with your own performance?), system satisfaction (How satisfied are you with the computer?), and comparative performance (How well do you think you perform compared to other participants?) on a 5-point Likert scale.

### EEG Recording & Preprocessing & Parameterization

EEG was recorded using 64 passive electrodes (Easycap GmbH, Herrsching, Germany) arranged according to the international 10–10 system and recorded with Neurone 1.5 recorder (Bittium Biosignals Ltd, Kuopio, Finland), and a NeurOne Tesla AC-amplifier (Bittium Biosignals Ltd, Kuopio, Finland). AFz served as the ground reference and FCz as online reference. Data was recorded with a sampling rate of 1000 Hz. Impedances were kept below 20 kΩ during recording.

EEG preprocessing used EEGLAB (Delorme & Makeig, 2004) in MATLAB. The online reference channel FCz was reconstructed by adding a placeholder channel and re-referencing the data to CPz. Subsequently, noisy channels were identified based on kurtosis and spectral criteria, were removed, and eventually interpolated using spherical splines (including CPz). Then, a common average reference was applied. Data were resampled to 200 Hz, band-pass filtered (2–30 Hz, 4th-order Butterworth), and epochs were created ranging from 0.8 s to 3 s relative to cue-onset. These epochs were then baseline corrected relative to the −0.2 s to 0 s period. After an automated detection and rejection of epochs of bad data quality, ICA was performed on PCA rank-compressed data. The resulting independent components (ICs) were classified via ICLabel (Pion-Tonachini et al., 2019). ICs with a probability of less than 0.30 to reflect brain activity, or more than 0.30 to reflect eye activity were removed from the data. For the cue-locked data M = 4.59 (SD = 2.29) channels, M = 132.27 (SD = 86.94) epochs, and M = 23.3 (SD = 6.14) ICs were removed. For the response/feedback-locked data, preprocessing was identical to the cue-locked data. Here, epochs were created ranging from −1.5 s to 1 s relative to the response/feedback onset. For the response locked data M = 4.59 (SD = 2.29) channels, M = 90.3 (SD = 64.25) epochs, and M = 22.14 (SD = 6.17) ICs were removed during preprocessing.

### TF decomposition and analysis

Cue-locked as well as response-locked EEG data was time–frequency decomposed using complex Morlet wavelet convolution on single-trial EEG data. For each participant, trials from the calibration block were excluded, and only trials with valid response times (RT < 1200ms) were retained. Remaining trials were assigned to their corresponding experimental condition (feedback type: veridical feedback, self-attributed errors, system-attributed errors) × (reward: high, low). Convolution was performed using 30 complex Morlet wavelets with logarithmically spaced centre frequencies between 2 and 30 Hz. The widths of the tapering Gaussians were also logarithmically spaced and chosen to yield temporal full-widths at half-maximum (FWHM) between 600 and 240ms. Wavelets were generated in the time domain, normalized to unit amplitude to ensure equal peak magnitude across frequencies, and transformed into the frequency domain for multiplication with the Fourier transform of the EEG data. After inverse fast-Fourier transformation, the complex analytic signal was obtained for each frequency and time point, from which single-trial power was extracted. Event-related spectral perturbations (ERSPs) were then computed by extracting single-trial spectral power and subsequently averaging power estimates within each condition, expressing these values as decibel change relative to a frequency-specific baseline ranging from −500 to −200ms relative to the time-locking event. Importantly, baseline power was computed across all trials independent of condition, which prevents condition-driven differences in the baseline interval from biasing normalization (cf. Arnau et al., 2020). Potential edge artifacts were controlled by pruning each epoch by 500ms at each side.

### Statistical analysis of EEG data

Time-Frequency representations of EEG activity were computed using EEGLAB for each participant and condition. We obtained ERSPs for all electrodes (condition × channel × frequency × time) and averaged across trials. For the statistical analysis, we obtained average ERSP values within a predefined region of interest (ROI) in sensor × frequency × time-space for each condition. The ROI for alpha power during the cue-target interval comprised the electrodes PO7 and PO8, the frequencies ranging from 8-12 Hz, and the time window from 1200-1500ms relative to cue-onset. For beta power the ROI were the electrodes C3, C4, CP3, and CP4, with the time-frequency window ranging from 16-30Hz and 1200-1500ms relative to cue-onset. For measures of response-related theta, the ROI comprised the electrodes FCz, Fz, FC1, FC2 and CZ, and the frequencies ranging from 4 to 7 Hz. For error-related activity, a time-window ranging from 100 to 400 ms relative to the response was used, for feedback-related activity, the time-window was 300 to 400 ms. Statistical testing was performed using RM-ANOVAs. For the analysis of occipital alpha power and central beta power during the cue-target interval, as well as for frontal midline theta related to error monitoring, a 3 (feedback type: veridical feedback, self-attributed errors, system-attributed errors) × 2 (reward: high, low) design was used. For the analysis of frontal midline theta related to flipped feedback, this design was reduced to 2 × 2 factors, as no feedback flips occurred in the veridical feedback condition. When significant main effects were observed, post-hoc pairwise comparisons were conducted using paired t-tests. Benjamini–Hochberg (BH) false-discovery rate (FDR) correction was applied to all post-hoc p-values. Standardized effect sizes (Cohen’s *d*) and Bayes Factors (BF₁₀) were computed for all pairwise contrasts.

### Statistical analysis of behavioral data

We focused our statistical analysis of behavior on response times (RT) and accuracy. To avoid confounding accuracy and variability, all RT analyses were performed on correct trials only. Furthermore, trials with a RT<1200 were excluded from all behavioral analyses. For each participant and condition, we calculated the mean RT (MRT), the variance of RT (VRT), and the proportion of correct responses (PC). We then conducted a 3 (feedback type: veridical feedback, self-attributed errors, system-attributed errors) × 2 (reward: high, low) within-subject RM-ANOVA on response times and on accuracy. When significant main effects or interactions were detected, post-hoc comparisons were performed using paired-sample t-tests with BH-correction for multiple comparisons.

Additionally, we decomposed behavioral performance into latent parameters, according to the drift diffusion model (Ratcliff, 1978). These are (1) drift rate (v), representing the rate at which evidence is accumulated, (2) boundary separation (a), indicating the amount of evidence required to make a decision and hence representing response caution, and (3) non-decision time (Ter), representing time spent on processes not linked to decision making, such as perceptual encoding or motor execution. These parameters of the diffusion model were estimated using a Python implementation of the EZ-DDM function (Wagenmakers et al., 2007), which uses mean response times and response times variance within each condition. The statistical analysis of these parameters was identical to the analysis of response times and response accuracy.

### Statistical Analysis of Thought Probes

To analyse the thought probes collected throughout the experiment, we grouped the data by participant and condition and calculated mean ratings for all items (focus, self-satisfaction, system-satisfaction, and performance). We then conducted separate RM-ANOVAs on each item to assess main effects of feedback type. We used pairwise comparisons with BH-correction to explore significant main effects.

## Results

### Behavioral performance

The results of the behavioral analyses are depicted in Figure 2. The analysis of response times showed a significant main effect of feedback type *F*(2, 68) = 8.34, *p* = .001, ηG² = .012. Post-hoc tests indicated that participants responded fastest in the veridical condition (*M* = 0.571s, *SD* = 0.061), significantly slower in self-attributed errors (*M* = 0.580s, *SD* = 0.056; *t*(34) = −2.09, *p* = .045, *d* = −0.16, BF₁₀ = 1.24), and slowest in system-attributed errors (*M* = 0.586s, *SD* = 0.055; vs. veridical: *t*(34) = −3.73, *p* = .002, *d* = −0.26, BF₁₀ = 43.3; vs. self: *t*(34) = −2.43, *p* = .030, *d* = −0.11, BF₁₀ = 2.35). A significant main effect could also be observed for the factor reward, *F*(1, 34) = 6.61, *p* = .015, ηG² = .010. Responses were faster in high (M = 0.577s, SD = 0.058) versus low reward trials (M = 0.581 s, SD = 0.057).

**Figure 2.**
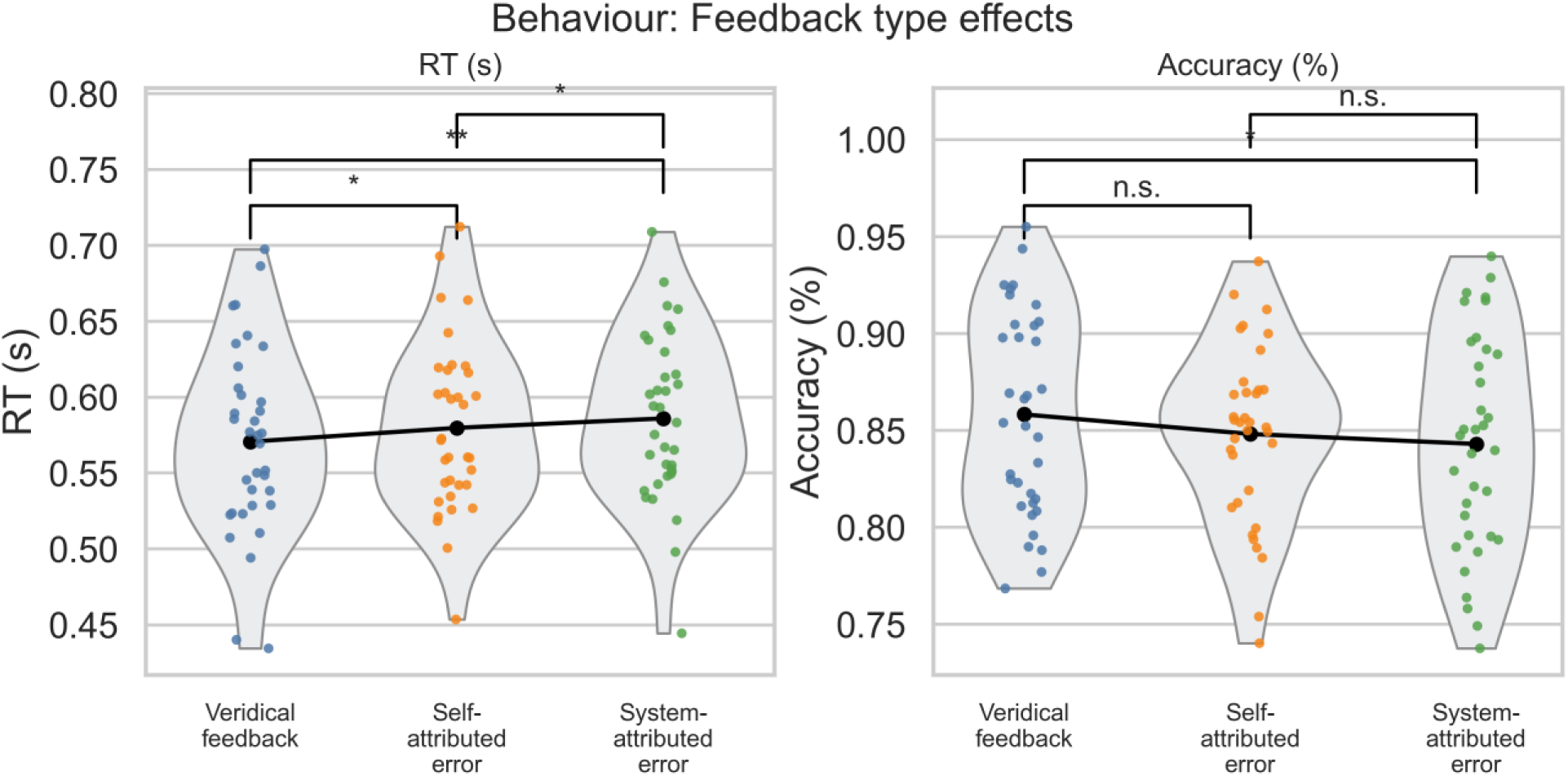
Behavioral performance as a function of feedback condition. Violin plots show distributions of mean response times (RT) and accuracy across the three feedback conditions (veridical/neutral, self-attributed errors, system-attributed errors). Black dots and connecting lines indicate condition means; brackets denote BH-corrected pairwise comparisons.

A significant main effect for the factor feedback type could also be observed for response accuracy (*F*(2, 68) = 4.38, *p* = .027, ηG² = .018). Post-hoc tests indicated higher accuracy for veridical than for system-attributed feedback (t(34) = 2.95, p = .017, d = 0.28, BF₁₀ = 6.98), whereas the comparisons between veridical and self-attributed feedback (p = .174) and between self- and system-attributed feedback (p = .289) were not significant.

### Drift-Diffusion Modelling (DDM)

Results from the EZ-DDM revealed that feedback type significantly modulated drift rate (v) and non-decision time (Ter), but not boundary separation (a). Drift rate differed across feedback types, F(2, 68) = 4.38, p = .027, with lower v for system-attributed than for veridical feedback (t(34) = 2.96, p = .020, d = 0.29, BF₁₀ = 5.98). Drift rate for self-attributed errors was intermediate (veridical vs. self: p = .100; self vs. system: p = .627, BF₁₀ < 1).

Non-decision time also differed by feedback type, F(2, 68) = 7.28, p = .002, with longer Ter for system-attributed than for both veridical feedback (t(34) = −3.43, p = .005, d = −0.25, BF₁₀ = 20.99) and self-attributed errors (t(34) = −2.60, p = .020, d = −0.14, BF₁₀ = 3.31). A significant main effect of reward was also observed, F(1, 34) = 5.86, p = .020, indicating that non-decision time was longer in low-reward than in high-reward trials. Boundary separation did not differ across conditions (all p > .15), indicating no evidence for changes in response caution. Overall, these results demonstrate that performance impairments under compromised agency reflect reduced rate of evidence accumulation (lower v) and slowed perceptual/motor processing (increased Ter), rather than increased decision caution (a).

**Figure 3.**
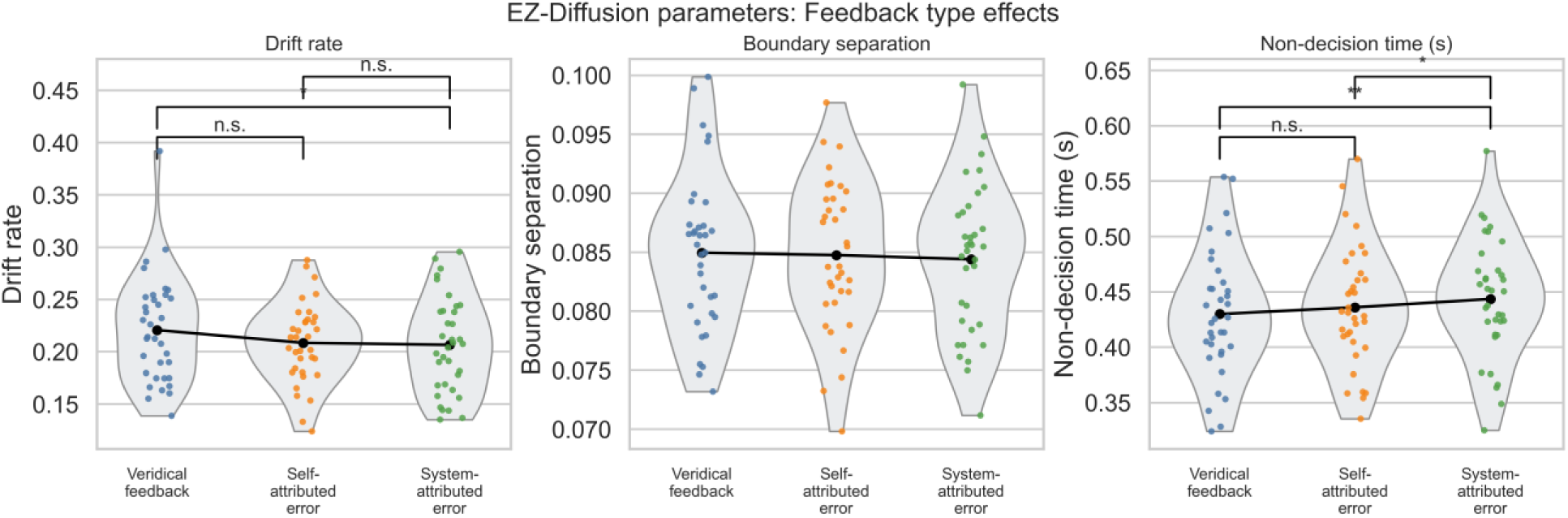
Diffusion parameters across feedback conditions. Violin plots show distributions of drift rate (v), boundary separation (a), and non-decision time (Ter) across the three feedback conditions (veridical/neutral, self-attributed errors, system-attributed errors). Black dots and connecting lines indicate condition means, brackets denote BH-corrected pairwise comparisons.

### Thought probes

Thought probes demonstrated a robust main effect of feedback type, *F*(2, 68) = 84.45, *p* < .001, ηG² = .246, no reward effect (*F*(1, 34) = 1.14, *p* = .293), and a modest interaction (*F*(2, 68) = 3.46, *p* = .047). Self-satisfaction was higher in the veridical feedback condition than in both self-attributed (*t*(34) = 9.10, *p* < .001, *d* = 1.81) and system-attributed conditions (*t*(34) = 8.70, *p* < .001, *d* = 1.66), with no difference between trials in the self-attributed and system-attributed error conditions (*p* = .580). Computer-satisfaction showed a graded decline: Computer satisfaction was higher under veridical feedback than under both self-attributed (t(34) = 6.75, p < .001, d = 0.81) and system-attributed feedback (t(34) = 6.55, p < .001, d = 1.16), and computer-satisfaction under self-attributed feedback exceeded system-attributed feedback (t(34) = 3.20, p = .003, d = 0.38). Focus was higher in the veridical feedback condition relative to both self-attributed and system-attributed error conditions (both p < .001), with no difference between self- and system-attributed errors (p = .522). The same pattern held for perceived performance, which was higher under veridical feedback than in the self- and system attributed error conditions (both p < .001), again with no significant self–system difference (p = .775).

**Figure 4.**
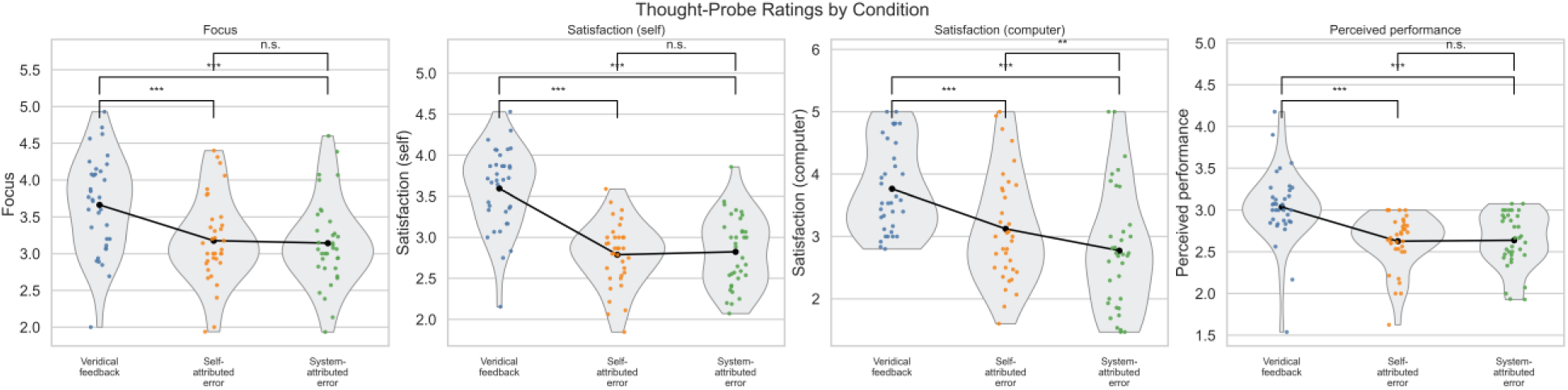
Thought Probe Ratings across Feedback Conditions. Violin plots show participant-level distributions for each rating (focus, self-satisfaction, computer-satisfaction, perceived performance) across veridical feedback, self-attributed error, and system-attributed error conditions. Dots represent individual participants’ mean ratings; black lines indicate condition means. Brackets show BH-corrected pairwise comparisons.

### EEG time–frequency analysis

Alpha power (1200–1500 ms; 8–12 Hz; PO7/PO8) in the cue-target-interval (CTI) showed a significant main effect of feedback type, *F*(2, 68) = 10.99, *p* < .001, ηG² = .015. Post-hoc tests revealed lower alpha power in veridical feedback trials than in both self-attributed (*t*(34) = –3.81, *p* < .001, *d* = –0.26, BF₁₀ = 53.9) and system-attributed trials (*t*(34) = – 4.43, *p* < .001, *d* = –0.27, BF₁₀ = 265.1), with no difference between error conditions. A main effect of reward was also found, *F*(1, 34) = 7.26, *p* = .011, ηG² = .004, reflecting lower alpha power for high- versus low-reward trials (M = –1.371 vs. –1.221; *t*(34) = –2.69, *p* = .011, *d* = –0.13, BF₁₀ = 3.97).

**Figure 5.**
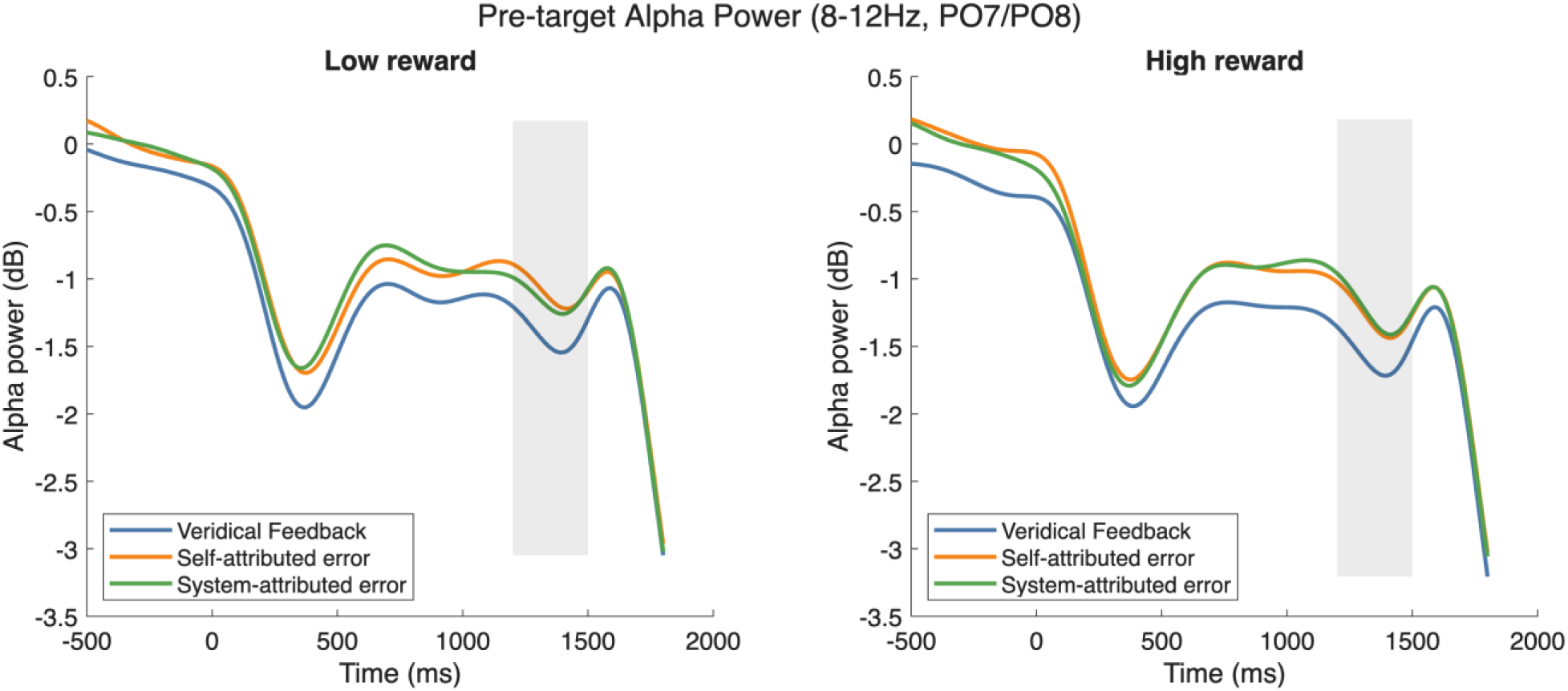
Pre-target alpha power in the CTI. Alpha band power (db) averaged over PO7/PO8 separated for low-(left) and high- (right) reward trials. Lineplots show the three feedback conditions (veridical feedback, self-attributed error, system-attributed error). The shaded region marks the predefined time window of analysis (1200-1500ms).

Beta power (1200–1500ms; 16–30 Hz; C3/C4/CP3/CP4) in the CTI showed significant main effects of feedback type, *F*(2, 68) = 3.74, *p* = .035, ηG² = .021, and reward, *F*(1, 34) = 5.28, *p* = .028, ηG² = .004, with no interaction. Post-hoc comparisons indicated that higher beta power was higher in self-attributed trials than in veridical trials (*t*(34) = –3.01, *p* = .015, *d* = –0.30, BF₁₀ = 7.95), whereas veridical versus system-attributed trials did not differ (*p* = .077). High-reward trials again showed lower beta power than low-reward trials (M = –0.782 vs. –0.722; *t*(34) = –2.30, *p* = .028, *d* = –0.14, BF₁₀ = 1.81).

**Figure 6.**
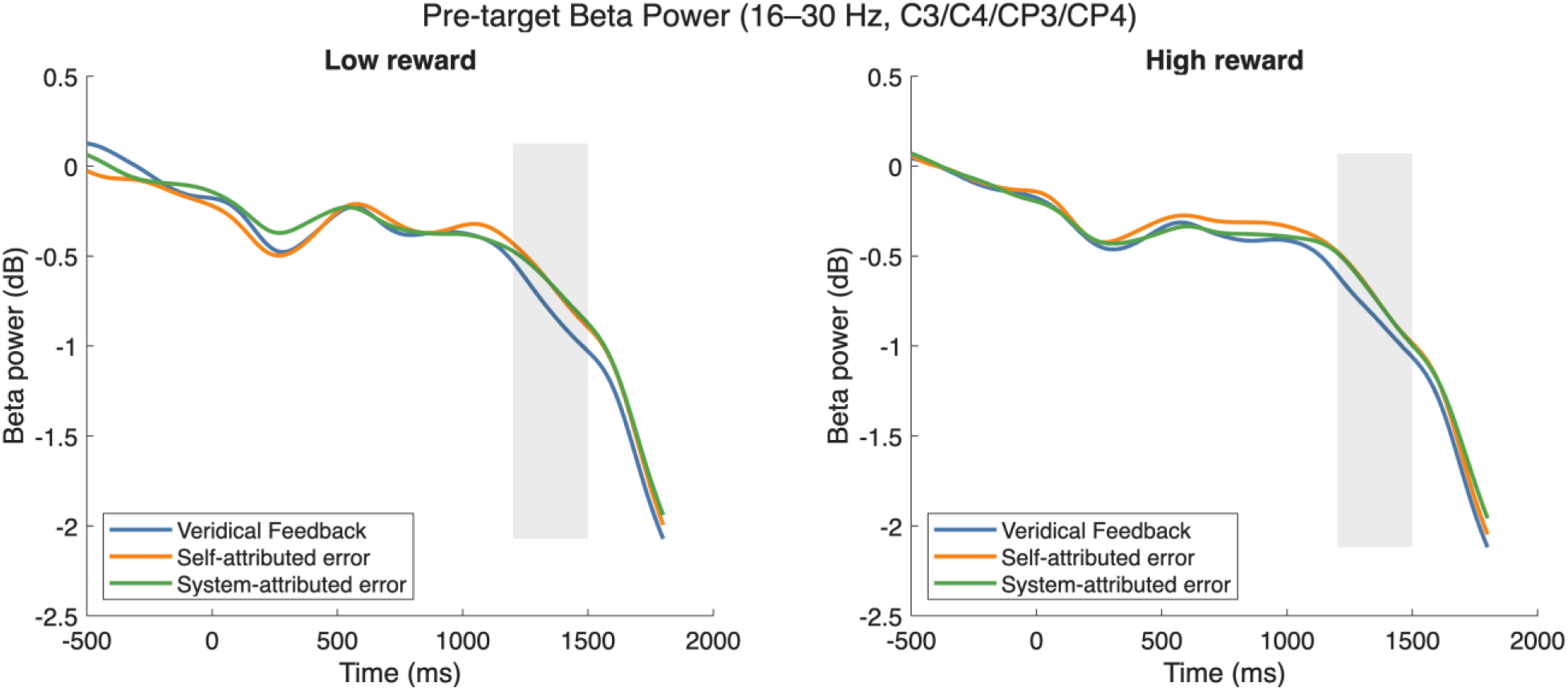
Pre-target beta power in the CTI. Beta band power (db) averaged over C3/C4/CP3/CP4 separated for low-(left) and high- (right) reward trials. Lineplots show the three feedback conditions (veridical feedback, self-attributed error, system-attributed error). The shaded region marks the predefined time window of analysis (1200-1500ms).

The difference in response-related mid-frontal theta power between error and correct trials (100–400 ms; 4–7 Hz; FCz/Fz/FC1/FC2/Cz) showed a significant main effect of feedback type, *F*(2, 68) = 14.83, *p* < .001, ηG² = .068 with no main effect of reward (*F* = .057, *p* = .456) and no interaction (*F* = 0.78, *p* = .461). Post-hoc comparisons revealed stronger theta power for veridical feedback than for system-attributed errors, t(34) = 4.24, p < .001, dz = 0.62, BF₁₀ = 164.22, and self-attributed errors, t(34) = 4.63, p < .001, dz = 0.61, BF₁₀ = 466.43. Theta power did not differ between system- and self-attributed error conditions, t(34) = −0.08, p = .936, dz = −0.01, BF₁₀ = 0.18.

**Figure 7.**
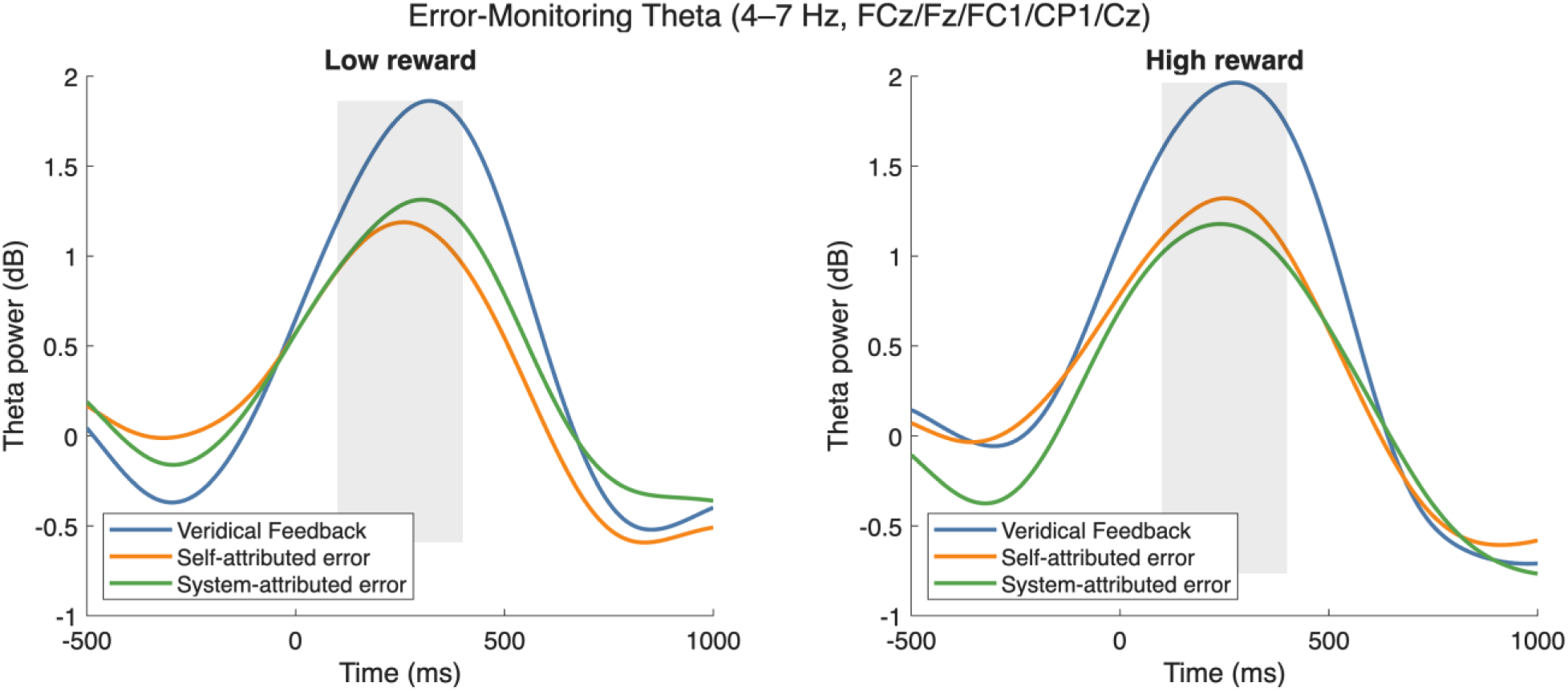
Response-related midfrontal theta-power during error monitoring. Theta band power (db) averaged over FCz/Fz/FC1/FC2/Cz separated for low-(left) and high- (right) reward trials. Lineplots show the three feedback conditions (veridical feedback, self-attributed error, system-attributed error). The shaded region marks the predefined time window of analysis (50-250ms post response).

Feedback-related frontal theta power (300-400ms; 4–7 Hz; FCz/Fz/FC1/FC2/Cz) on self-and system-attributed trials with flipped feedback showed a significant main effect of feedback type, with stronger theta responses for system-attributed error than for self-attributed errors, *F*(1,34) = 13.50, *p* < .001, ηG² = .036.

**Figure 8.**
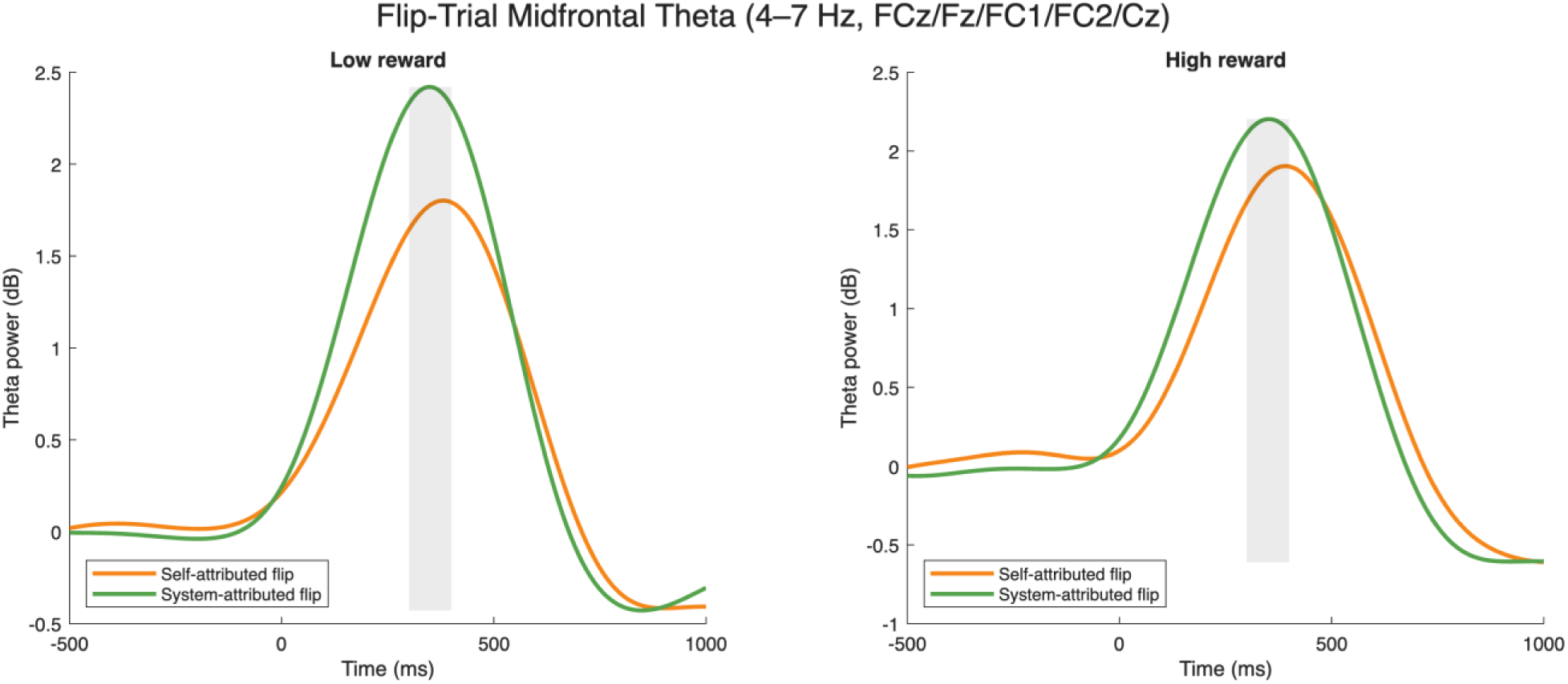
Feedback-locked midfrontal theta-power during monitoring of flipped feedback. Theta band power (db) averaged over FCz/Fz/FC1/FC2/Cz separated for low-(left) and high- (right) reward trials. Lineplots show two feedback conditions (self-attributed error, system-attributed error). The shaded region marks the predefined time window of analysis (300-400ms post response).

## Discussion

The present study examines the effect of sense of agency (SoA) on task engagement, with particular emphasis on the role of error attribution. To this end, we implemented experimental conditions involving manipulated performance feedback. These manipulations targeted (1) SoA directly, by presenting correct responses as incorrect, and (2) error attribution, by providing feedback that either indicated the participant’s response was incorrect or suggested that an unpressed button had been registered as pressed. Importantly, this manipulation was structured in a block-wise manner, with block transitions occurring in the background and not signaled to the participants. In this way, we obtained episodes of veridical feedback, episodes of feedback suggesting increased error rates due to one’s own mistakes (self-attribution), and episodes of feedback suggesting increased error rates due to system failure (system-attribution).

Thought probes collected throughout the experiment indicate that the manipulation worked as intended. Participants reported significantly higher levels of satisfaction with their own performance in the veridical feedback condition than in the self- and system-attribution conditions, with no difference between the latter two. Regarding satisfaction with system performance, however, the difference between the self- and system-attribution conditions was significant. We also manipulated reward on a trial-by-trial basis to investigate whether the effects of the feedback manipulation differed between high- and low-reward trials.

The behavioral results are largely in line with our expectations. Participants responded fastest in the veridical feedback condition, significantly slower in the self-attribution of errors condition, and even slower in the system-attribution of errors condition. As trial difficulty was fixed for each participant, this pattern suggests that it is not only the likelihood of successful task performance that governs cognitive engagement, but also critically how failure is attributed (Synofzik et al., 2008; Homsma et al., 2007). Decomposing response times into parameters of the drift diffusion model using the EZ-diffusion algorithm indicates that the response time differences may be driven by two underlying mechanisms. Regarding drift rate, the parameter assumed to reflect the speed of evidence accumulation, significant differences were observed between the veridical feedback and system-attribution conditions. Non-decision time, which captures variance related to motor execution and sensory encoding, was significantly larger in the system-attribution condition than in both other conditions. Importantly, boundary separation remained unchanged, arguing against a speed–accuracy trade-off. Together, these findings indicate that compromised controllability reduces efficiency of information processing, with additional behavioral costs when action–outcome contingencies are externally disrupted.

In the EEG analyses, we investigated posterior alpha power and centro-parietal beta power during the cue–target interval as measures of proactive cognitive control. Posterior alpha power has been associated with cortical excitability and visuospatial attention (Jensen & Mazaheri, 2010; Klimesch, 2012; Foxe & Snyder, 2011). Prior to an imperative stimulus, it is typically interpreted as a correlate of attentional engagement, with stronger task orientation reflected in greater decreases in alpha power (Cooper et al., 2003; Hanslmayr et al., 2011). Alpha power has been shown to index modulations of attentional engagement across a wide range of research domains, including mental fatigue (Arnau et al., 2021; Tran et al., 2020; Wascher et al., 2014), reward prospect (Arnau et al., 2024; Sawaki et al., 2015; van den Berg, 2014), mind wandering (Arnau et al., 2020; Compton et al., 2019), and task load (Jensen et al., 2002; Klimesch, 2012). Similarly, centro-parietal beta power during task preparation has been shown to scale with motor readiness (Pfurtscheller & Lopes da Silva, 1999). Elevated beta activity has been linked to the maintenance of existing sensorimotor states and reduced initiation of forthcoming actions (Engel & Fries, 2010; Luft et al., 2013). Moreover, Bu-Omer et al. (2021) demonstrated that parieto-occipital alpha and low beta-EEG power increased with reduced subjective SoA, suggesting that these frequency bands are sensitive markers of agency disruptions.

In line with this, alpha and beta power in the present study were significantly higher during the cue–target interval when feedback indicated a higher error rate. In this way, our results indicate that the manipulated feedback affected proactive control processes, which is consistent with the theoretical assumption that a higher feedback-induced error rate reduces SoA on a global level. Within control allocation frameworks, perceived controllability is a critical determinant of expected control efficacy. Therefore, a reduced SoA should in turn lead to reduced proactive cognitive effort, as the likelihood of obtaining reward is diminished (Chiew & Braver, 2014; Fröber & Dreisbach, 2016). Consequently, the subjective value of exerting effort decreases (Shenhav et al., 2013; Shenhav et al., 2017). Previous work has shown that both prospective measures and retrospective outcome congruency independently modulate implicit and explicit agency measures, underscoring that agency arises from multiple cue integrations (Barlas & Kopp, 2018).

Taken together, these preparatory oscillatory effects reflect cognitive states associated with perceived action–outcome contingencies: When reliability decreases, task preparation decreases accordingly. Importantly, these preparatory effects in the alpha and beta bands did not differentiate between the self- and system-attribution conditions. This suggests that proactive control allocation is sensitive to overall controllability rather than the experienced source of failure (Mittelstädt et al., 2024). It further indicates that variance in these neural measures cannot fully account for the observed behavioral differences.

In addition to the cue-locked analysis of preparatory processes during the cue-target interval, we also investigated effects in the response-related data. As the feedback was presented with a zero-latency relative to the response, this data is feedback-locked as well. The focus of our analysis was error monitoring and feedback evaluation. To this end, we focused on time-frequency power in the theta band at mid-frontal electrodes. Midfrontal theta has been widely implicated in signaling the need for control when conflicts or errors arise (Cavanagh & Frank, 2014; Cavanagh & Shackman, 2015; Helfrich & Knight, 2016). Theta dynamics further clarify how the monitoring system adapts to differences in feedback reliability and error source. The response-related theta response was strongest following veridical outcomes and reduced for both self- and system-attributed errors, consistent with models in which reliable action-outcome mappings enhance the salience of performance-relevant deviations (Cavanagh & Frank, 2014; Cohen, 2014). In contrast, during trials with manipulated feedback, the outcome explicitly conflicted with the participant’s action. In these trials, midfrontal theta increased more for system- than self-attributed errors, reflecting stronger action-outcome conflict and the engagement of attributional updating processes (Cavanagh & Shackman, 2015). Critically, our data indicate that such theta activity indicates not only the presence of conflict, but also its attributional context. When errors were externally attributed, theta increases likely reflected a violation of the action-outcome-contingency itself. In contract, self-attributed errors were processed within a preserved agency framework, resulting in different evaluative demands. This interpretation is consistent with recent evidence that the temporal unfolding of neural oscillations predicts agency judgments (Veillette et al., 2023), supporting accounts in which SoA emerges from dynamic integration of pre- and post-movement signals. Together, these findings suggest that theta power does not simply track errors but is also sensitive to the inferred source of action-outcome-discrepancies. It differentiates whether an action-outcome conflict reflects an impairment of the own performance under intact causal structure, or a disruption of the action-outcome contingency itself.

Although we observed small additive reward effects on RT and preparatory oscillations, these were not modulated by agency reliability and did not interact with control allocation. This suggests that perceived outcome controllability, rather than incentive value, was the primary determinant of preparatory investment. This aligns with control allocation models (Shenhav et al., 2013), which predict that expected efficacy of control outweighs reward magnitude when allocating cognitive effort.

Our findings can be related to the classic comparator model (Wolpert et al., 1995; Blakemore et al., 2000), which proposes that SoA arises from comparing predicted sensory consequences of an action with actual outcomes. Stronger alpha/beta desynchronization under veridical feedback fits this framework as evidence of robust predictive signaling, while enhanced theta under errors reflects postdictive mismatch detection. While this captures important elements in our data, the results also indicate that this mechanism alone cannot account for context-dependent cue integration, motivating broader inference-based models. A pure prediction-outcome comparator mechanism does not suffice to explain why externally attributed errors elicit stronger monitoring signals than self-attributed errors despite comparable preparatory signals. This points towards the necessity of source inferencing beyond mismatch detection. Extended comparator models emphasize prediction-outcome matching, in which preparatory oscillations (alpha/beta) reflect predictive signaling of motor intentions, while feedback-locked theta indexes postdictive comparison of predicted versus actual outcomes (Wen & Haggard, 2020). Our findings are consistent with this view: attenuated preparatory desynchronization under compromised agency illustrates how disrupted predictive cues (e.g., corrupted efference copies) reduce SoA. However, recent work has shown that the temporal dynamics of oscillatory signals can themselves predict agency judgments (Veillette et al., 2023), underscoring the need for inference-based accounts that move beyond pure prediction-outcome matching. Such multifactorial models emphasize the role of cue integration (Synofzik et al., 2008, 2013). Gentsch et al. (2012) have demonstrated that SoA judgements flexibly draw on motor-and non-motor anticipatory cues given their relative reliability. Our theta results fit this perspective: context-dependent shifts (error salience vs. conflict) reflect adaptive integration of predictive and postdictive feedback, consistent with predictive coding frameworks (Friston, 2012). Thus, agency computations in our task emerge from a continuous interplay between preparatory predictive signals and postdictive feedback evaluation. Building on these descriptive accounts, Bayesian computational models provide a unifying framework in which comparator-derived prediction errors and contextual priors are weighted by precision to yield SoA (Legaspi & Toyoizumi, 2019; Tanaka, 2024; Dutta, 2025). In our task, compromised feedback reduced the expected precision of action-outcome mappings, leading to diminished preparatory investment. However, the attribution of errors to oneself versus the system selectively altered the inferred source of prediction errors, thereby shaping monitoring signals without equivalently affecting proactive control allocation. This is consistent with active inference accounts, in which disengagement reflects an optimal strategy: when action-outcome priors are contradicted, reduced preparatory investment minimizes metabolic costs (Friston, 2010). Our results therefore illustrate how comparator mechanisms are embedded within a broader Bayesian inference process, where SoA dynamically shapes cognitive control. These findings suggest that agency-related neural dynamics dissociate along two dimensions: preparatory oscillations track perceived controllability of action-outcome contingencies, whereas midfrontal theta encodes the attribution-specific conflict that differentiate self-generates errors from externally generated errors. Together, these results position error attribution as a central component in the agency inference process, highlighting how prediction errors are interpreted and integrated without necessarily altering proactive control allocation.

## Conclusion

Our findings demonstrate functionally dissociable components within a recursive agency inference loop. Perceived outcome controllability primarily shaped proactive control allocation: reliable action-outcome mappings under veridical feedback were associated with stronger reductions in preparatory alpha and beta power, increased speed of evidence accumulation, and more efficient response behavior. In contrast, disrupted action-outcome controllability attenuated these preparatory dynamics on a global level. Oscillatory activity related to preparatory processes therefore indicate a current estimate of action-outcome controllability. However, error attribution selectively modulated outcome evaluation: when discrepancies were attributed to the system, midfrontal theta responses and behavioral costs amplified, suggesting heightened attribution-specific conflict. In this way, midfrontal theta encodes precision-weighted predication errors and their attributional interpretation. Within a Bayesian framework, our results suggest that SoA operates at two computational levels that feed information back into each other: it regulates the expected valence of action-outcomes during task preparation, and shapes how predication errors are attributed and integrated during outcome monitoring. In this way, comparator-derived mismatch signals are embedded within a broader inferential structure in which perceived controllability and attributional source inference determine the allocation of cognitive effort.

## Acknowledgements

We thank Tobias Blanke for his technical support of the experiment, and Nathalie Hutterloh, Oliver Butzke and Quang Dang for their efforts in data collection.

## Open Practice Statement

The data and code for all analyses will be made available before journal submission.

